# Reconstructing the evolutionary history of a B cell lineage with minimum spanning tree and genotype abundances

**DOI:** 10.1101/2022.02.27.481992

**Authors:** Nika Abdollahi, Lucile Jeusset, Anne de Septenville, Frédéric Davi, Juliana S. Bernardes

## Abstract

B cell receptor (BCR) genes exposed to an antigen undergo somatic hypermutations and Darwinian antigen selection, generating a large BCR-antibody diversity. This process, known as B cell affinity maturation, increases antibody affinity, forming a specific B cell lineage that includes the unmutated ancestor and mutated variants. In a B cell lineage, cells with a higher antigen affinity will undergo clonal expansion, while those with a lower affinity will not proliferate and probably be eliminated. Therefore, cellular (genotype) abundance provides a valuable perspective on the ongoing evolutionary process. Phylogenetic tree inference is often used to reconstruct B cell lineage trees and represents the evolutionary dynamic of BCR affinity maturation. However, such methods should process B cell population data derived from experimental sampling that might contain different cellular abundances. There are a few phylogenetic methods for reconstructing the evolutionary history of B cell lineages; best-performing solutions are time-demanding and restricted to analyzing a reduced number of BCR IGH sequences, while time-efficient methods do not consider cellular abundances. We propose ClonalTree, a low-complexity and accurate approach to reconstruct B cell lineage trees that incorporates genotype abundances into minimum spanning tree (MST) algorithms. Using both simulated and experimental data, we demonstrated that ClonalTree outperforms MST-based algorithms and achieves a similar performance compared to a method that explores tree generating space exhaustively. However, ClonalTree has a lower running time, being more convenient for reconstructing phylogenetic lineage trees from high-throughput BCR sequencing data, mainly in biomedical applications, where a lower computational time is appreciable. It is hundreds to thousands of times faster than exhaustive approaches, enabling the analysis of a large set of sequences within minutes or seconds and without loss of accuracy. The source code is freely available at github.com/julibinho/ClonalTree.

## 1 Introduction

B cells are essential components of the adaptive immune system. They express a cell surface receptor, the B cell receptor (BCR), recognizing a vast array of antigens. The main components of the BCR are immunoglobulins (IG), which can be secreted in a soluble form as antibodies. IGs are heterodimers composed of two identical heavy chains (IGH) and two identical light chains (IGL); each chain possesses a variable and a constant region. The variable regions are responsible for antigen-binding specificities, whereas the constant regions are associated with cell-signaling components of the BCR and facilitate interactions with other immune-system molecules. The variable regions are encoded by Variable (V), Diversity (D) in the case of heavy chains, and Joining (J) genes, which are rearranged during lymphopoiesis by a complex genetic mechanism known as V(D)J recombination (*1*). During this process, the combinatorial diversity is generated by ligating these genes into an exon, further enhanced by random deletions and insertion at their joining sites. These mechanisms allow newly-formed naive B cells to express a vast repertoire of distinct BCRs (> 10^13^) (*2*).

When exposed to an antigen, naive B cells’ genes encoding the IG undergo multiple rounds of mutations, e.g., somatic hypermutations (SHM) and Darwinian antigen selection. This stage of the B cell development is known as affinity maturation since it leads to a progressive increase of the IG’s affinity for their cognate antigens and occurs in specialized structures of secondary lymphoid organs, the germinal centers. During affinity maturation, B cells encounter numerous biological processes such as antigen presentation, proliferation, and differentiation. Natural selection selects B-cells with higher IG-antigen affinities, which will proliferate and undergo clonal expansion, while those with lower affinity will be eliminated (*3*). Affinity maturation produces a functionally heterogeneous population with different B cell lineages, each formed by the naive B cell and its variants. Thus, the number of unique variant sequences and their respective abundances (genotypes) provides an important perspective on the ongoing evolutionary process and helps elucidate clonal selection.

Understanding BCR repertoire evolution is necessary to answer fundamental biological questions such as clonal selection during antigen challenges, immune senescence, development of efficient vaccines, therapeutic monoclonal antibodies, or further understanding B cell tumor development. Recently, an evolutionary approach has been able to quantify dissimilarity when comparing BCR repertoires of young and aged individuals after influenza vaccination (*4*). Antibody evolutionary studies have also been used to guide the clonal reconstruction of BCR repertoires (*5, 6*).

Several studies have analyzed B cell clonal evolution (B cell lineage trees) to understand the evolutionary mechanisms involved in several diseases (*7, 8*). In this later contribution, the authors showed that lineage trees are largely shaped by antigen-driven selection occurring during an immune response. Genetic evolution, such as observed during affinity maturation, is often modeled through phylogenetic inference, a well-known methodology that describes the evolution of related DNA or protein sequences in various species. Theoretically, phylogenetic inference methods could reconstruct B cell lineage trees by replacing species with BCR IGH sequences (with different mutations). However, in a phylogenetic tree, the root is usually unknown, the observed sequences are usually represented only in the leaves, and the inner nodes represent the relationships amongst sequences. Conversely, in a B cell lineage tree, the root or the BCR IGH sequence of the naive B cell giving rise to the lineage can be deduced by aligning the IG variable region sequences with sequences stored in a reference database for identifying the germline V(D)J genes from which they derive (*9*). Another distinct point is that B cells with different BCR mutations can coexist; therefore, the observed BCR IGH sequences can be leaves or internal nodes in the tree (*10, 11*). Due to simultaneous divergence, multifurcations are also common (*12*). IG sequences are under intense selective pressure, and the neutral evolution assumption is invalid. Moreover, the context dependence of SHM violates the assumption that sites evolve independently and identically. Under these circumstances, conventional phylogenetic tree algorithms seem unsuitable for reconstructing B cell lineage trees. The performance of such methods varies substantially in terms of the tree topology and the ancestral sequence, as shown previously (*12, 13*).

Some computational tools were explicitly designed to reconstruct B cell lineage trees. Igtree (*14*) employs the maximum parsimony criterion to find the minimal set of events that could justify the observed sequences. It first constructs a preliminary tree, which only contains observed sequences, then uses a combined score based mainly on sequence mutations to gradually add internal nodes (unobserved sequences). IgPhyML combines the maximum likelihood approach (*15*) with a codon substitution model that uses a Markov process to describe substitutions between codons (*16*). IgPhyML has modified the codon substitution model to incorporate hot/cold-spot biases observed in BCR IGH sequences (*17*). GCtree (*18*) employs the maximum parsimony principle and incorporates the cellular abundance of a given genotype in phylogenetic inference. This information is used for ranking parsimonious trees, obtained by dnapars (*19*) with the assumption that more abundant parents are more likely to generate mutant descendants. GCtree uses a likelihood function based on the Galton-Watson Branching process (*20*). It is an accurate method, but its computational complexity is high, especially for a high number of sequences. GlaMST (*21*) is another method for reconstructing B cell lineage trees. It is a minimum spanning tree (MST)-based algorithm and iteratively grows the lineage tree from the root to leaves by adding minimal edge costs. GLaMST is more time-efficient than GCtree, but it ignores genotype abundance information.

Here we propose ClonalTree, a method to reconstruct B cell lineage trees, combining MST and genotype abundance to infer maximum parsimony trees. ClonalTree starts from the root (the inferred ancestral sequence) and iteratively adds nodes to the tree presenting minimal edge cost and maximum genotype abundance; therefore, it optimizes a multi-objective function rather than a single function based only on edge costs as implemented in GLaMST. Using simulated and experimental data, we demonstrate that ClonalTree outperforms GLaMST and achieves a comparable performance to GCtree. ClonalTree has a lower time complexity and great potential for many applications, particularly in clinical settings where time constraint is important.

## 2 Approach

ClonalTree reconstructs B cell lineage trees based on the minimum spanning tree (MST) and cellular (genotypes) abundances. We start with a formal description of the B cell lineage tree reconstruction problem, BCR IGH sequence distance, and minimum spanning tree algorithms. Next, we describe how we modified Prim’s algorithm (*22*) to incorporate genotype abundance information. Ultimately, we show how trees can be improved by creating intermediate nodes that describe non-observed sequences or performing editing operations.

### 2.1 Problem statement

Given a set of observed BCR IGH sequences with the same V(D)J rearrangement event and an inferred naive/unmutated BCR IGH sequence, we look for a minimum-sized directed tree structure: the B-cell lineage tree, which represents the affinity maturation process. Vertices (nodes) represent BCR IGH sequences, and the weight of edges connecting vertices represents the distance between sequences in terms of mutation, insertion, and deletion operations. All observed sequences are reachable from the root, the inferred naive/unmutated BCR IGH sequence.

### 2.2 Minimum spanning Tree

Given a connected, undirected graph (V, E), where V is a set of vertices, and E is the weight edges, its minimum spanning tree (MST) is a subset of vertices and edges that form a tree (a connected graph without cycles/loops) so that the sum of the weights of all the edges, the cost, is at minimum. For a given connected graph, several MSTs can exist. All trees have similar costs, but their topologies are different. Most MST construction algorithms are greedy approaches, where edges are sorted according to their weights and selected according to some criteria. In each step, greedy algorithms make a locally-optimal choice hoping that this choice will lead to a globally optimal solution.

### 2.3 A modified Prim’s algorithm

Prim’s (*22*) and Kruskal’s (*23*) are algorithms for finding the minimum spanning tree of a graph. Both are greedy approaches and present low time complexity. However, Prim’s algorithm runs faster than Kruskal in dense graphs (*24*). Therefore, we modified Prim’s algorithm to reconstruct B cell lineage trees. We start at the root and add all its neighbors with minimum edge weight to a priority queue. We then iteratively extract from the priority queue the node with the lowest edge weight and highest genotype abundance. If no cycle is formed, the node and the edge will be added to the tree. For each added node, all its neighbors with minimum edge weights are included in the priority queue. We keep on adding nodes and edges until we cover all nodes. In order to decrease the time complexity of the algorithm, we add each node only once to the priority queue. The original Prim’s algorithm has only one objective function, which minimizes the sum of edge weights (cost). Here we include a second objective function to maximize genotype abundance. If a set of edges have the same weight, we will choose the one that connects nodes with high abundance. Prim’s algorithm has a time complexity of *O*(|*V* |^2^) in the worst case but can be improved up to *O*(|*E*| + log |*V* |) when using data structures based on Fibonacci heaps (*25*). Figure 1 shows a simple example of the tree construction process.

**Figure 1:**
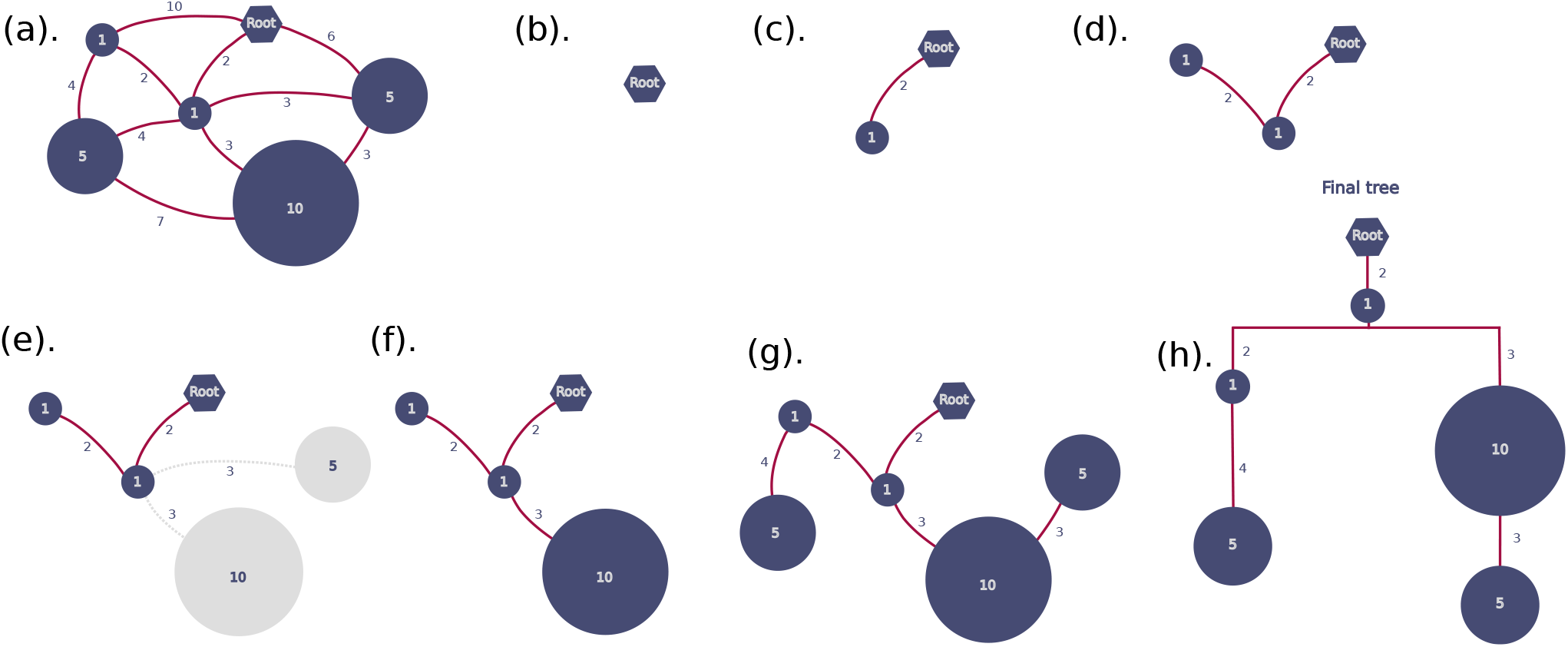
ClonalTree construction example. We start with a connected weighted graph (a) where nodes represent BCR genotype sequences, edge weights their distances, and node weights their abundances. The graph can be fully connected, or one can disable edges whose weight is lower than a threshold *δ*. Then, we first place the inferred ancestral sequence (the root) (b) and iteratively add nodes to the tree with the lowest edge weight and highest genotype abundance (c,d); when edges have the same weight (e), we choose that connected to the node with the highest abundance (f), we repeat until all nodes were added to the tree (g), the final tree is shown in (h).

### 2.4 Measuring genotype abundances

A B cell lineage tree might represent the relationships between the naive BCR IGH sequence and its mutants. Multiple copies of a given variant could indicate its importance within the affinity maturation and clonal expansion processes. Thus, we used variant abundances (here called genotype abundance) to guide the B cell lineage tree reconstruction. A genotype can be defined in different ways. It can represent a set of identical sequences or a group of similar sequences, where some mutations are allowed. A common way of defining a BCR genotype is to group sequences with the same V and J genes and the identical CDR3 amino acid sequences. In both cases, the genotype abundance accounts for the number of such elements within a populations of BCR IGH sequences.

### 2.5 Computing BCR IGH sequence distances

The reconstruction of B cell lineage trees requires a distance measurement between clonally-related sequences. This distance is usually used to weigh edges in the graph. Several works have used pairwise edit distances, such as Levenshtein (*26*), but its computation usually requires dynamic programming algorithms that become computationally expensive in time and memory when the number of sequences per lineage tree is large. Another weak point of pairwise distances is that they provide a local landscape without considering mutated regions across all the sequences. ClonalTree first performs a multiple sequence alignment to consider evolutionary events such as mutations, insertions, and deletions over all clonally-related sequences. Next, it computes a normalized hamming distance (*27*) between each pair of genotype sequences and then uses it as the weight edge connecting two genotypes. In order to avoid a fully connected graph, we can apply a threshold *δ*, that will disable edges whose weight is lower than *δ*.

### 2.6 Editing the reconstructed lineage tree

A greedy algorithm makes the optimal choice at each step, attempting to find the optimal way to solve the optimization problem. It never reconsiders its choices, while optimal algorithms always find the best solution. A way to review decisions and improve the ClonalTree algorithm is to edit the obtained lineage tree. Thus, we implemented two strategies: add unobserved intermediate nodes to the tree and detach/reattach sub-trees.

Unobserved internal nodes might represent unobserved sequences that were not sampled or disappeared during affinity maturation. In those cases, the evolutionary relationships were also lost. One way to recover them is to analyze the reconstructed tree to identify common ancestors not yet represented. This process is similar to building a phylogenetic tree among species, where unobserved internal nodes represent common ancestors of descendants. However, only leaves’ nodes are observed in a classical phylogenetic tree, and all internal nodes are unobserved, while in a B cell lineage tree, internal nodes can be observed or unobserved. When we detect a missing common ancestral in the tree, we add an unobserved internal node. It generally happens when we observe a distance between sister nodes that is smaller or equal to the distance for their parent, see nodes {d, e} in Figure S1-A. To add unobserved internal nodes, we traverse the tree in a pre-order manner, and for each pair of sister nodes, we verify if a common ancestor is missing. If it is the case, we add an unobserved internal node in the tree, connect it to the observed nodes by direct edges Figure S1-B, and update edge weights Figure S1-C. For this later, let *d*_*pm*_ and *d*_*pn*_ be the edge weights connecting a pair of sister nodes *m* and *n* to their parent *p*, respectively. Let *d*_*mn*_ be the distance between the sister nodes and *i* an unobserved internal node added to the tree. The edge weights are updated as follows:

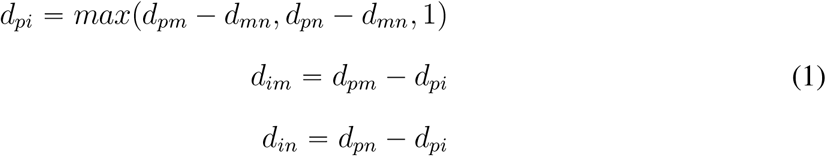

We can detach a sub-tree from an internal node by removing its edge and reattaching it to another internal node or leaf. We perform this editing operation to reduce the depth of the lineage tree by keeping the overall cost. The motivation here is to try to find the most parsimonious tree. We consider all branching nodes (i.e., nodes with more than one descendent) for this edition operation. Then, we try to detach and reattach each node (under edition condition) to any other node in the lineage tree. If this operation reduces the tree depth, we accept it and examine the resulting lineage tree again for additional edition operations that may further reduce the tree depth. We repeat this process until no editing operation can reduce the depth of the tree (see an illustration in Figure S2).

## 3 Methods

In the following, we describe how data sets were obtained or produced, how the performance of B cell lineage tree algorithms was evaluated, and the tools considered in inter-comparisons.

### 3.1 Data sets

We used two types of data sets to measure the ClonalTree performance and compare it with existing algorithms: simulated and experimental. GCtree simulator produced simulated data (*18*), while one of two experimental data sets was created during chronic lymphocytic leukemia (CLL) routine diagnostic procedures at the Pitié Salpêtrière hospital (Paris-France), and the second one is a public data set (*28*).

#### Simulated lineage trees

In order to create simulated lineage trees, we used the B cell lineage simulator provided by GCtree. The simulator produces a B cell lineage by randomly selecting IGHV, IGHD, and IGHJ germline genes from the IMGT database. Then, nucleotide(s) can be added to or removed from the junction region: IGHV-IGHD and IGHD-IGHJ. Next, it performs a branching process, and point mutations can be included in the descendants. Somatic hypermutations are simulated by a sequence-dependent process, where mutations are preferentially introduced within specific hotspot motifs (*29*). We kept simulator default parameters and generated 92 artificial lineage trees. The sizes of simulated trees ranged from 6 to 99 nodes, the number of observed sequences between 20 and 200, the degree of root nodes varied from 1 to 42, and depth trees from 2 to 7, see Table 1.

**Table 1:** Characteristics of simulated lineage trees.

|  | min | mean | max | std |
| --- | --- | --- | --- | --- |
| Tree depth | 2 | 4 | 7 | 1 |
| Degree of Root | 1 | 11 | 42 | 9 |
| Number of nodes | 6 | 34 | 99 | 21 |
| Number of leaves | 5 | 25 | 82 | 16 |
| Number of observed sequences | 30 | 81 | 200 | 48 |

##### Experimental data

We used a public data set from a previously reported experiment (*28*), where the authors combined multiphoton microscopy and single-cell mRNA sequencing to obtain IGH sequences extracted from germinal B cells of a lineage sorted from the mouse germinal center. The data set, labeled as TAS-42, contains 65 IGHV sequences and 42 genotypes; it is available at https://github.com/matsengrp/GCtree/tree/master/example. For this data set, a genotype is a set of identical sequences. The ancestor IGHV gene was inferred with Partis (*30*), and the final data set contains 66 sequences: 65 from the original data set plus one representing the reconstructed germline sequence.

The second experimental data set was generated by sampling sequences from the most abundant clone of a BCR repertoire associated with a CLL patient. The data set contains 3406 sequences, annotated with IGHV4-34*01/IGHD3-3*01/IGHJ4*02 genes using IMGT/HighV-QUEST software (*31*). For this data set, a genotype groups identical BCR IGH sequences; we obtained 20 genotypes with different abundances. Thus, the data set was labeled as CLL-20. To predict the hypothetical naive sequence (the root of this lineage tree), we considered the germline sequences of the corresponding IGHV and IGHJ genes provided by IMGT/HighV-QUEST. For the junction, we took it from the sequence with the lowest number of mutations on the IGHV gene when compared to the germline determined by IMGT/HighV-Quest. Eventually, we concatenate germline IGHV, the junction sequence, and the IGHJ sequences to obtain the hypothetical naive sequence.

### 3.2 Tree comparison and evaluation

To evaluate the performance of B cell lineage reconstruction algorithms, we used two metrics to compare tree topologies: graph editing distances (*32*), and ancestral sequence inferences (*12, 18*).

#### Graph Editing Distance

Let G_1_ and G_2_ be two graphs; the Graph Editing Distance (GED) finds the minimum set of graph transformations able to transform G_1_ into G_2_ through edit operations on G_1_. A graph transformation is any operation that modifies the graph: insertion, deletion, and substitutions of vertices or edges. GED is similar to string edit distances such as Levenshtein distance (*26*) when we replace strings by connected directed acyclic graphs of maximum degree one. We used two versions of GED, one applied to the whole tree (GED tree-based), and another applied to each branch separately (GED path-based). The latter version is more stringent than the first one since any difference in the path from each leaf to the root is considered a transformation.

The problem of computing graph edit distance is NP-complete (*33*), and there is no optimal solution in a reasonable time. This problem is difficult to approximate, and most approximation algorithms have cubic computational time (*34, 35*). Here we could use an optimal algorithm implementation since the number of nodes of evaluated trees was small. Nevertheless, we favored a grid cluster to compute GED for trees with more than 50 nodes. The code is available at https://github.com/julibinho/ClonalTree.

##### Ancestral reconstruction distances

We also compared trees by measuring their ancestral sequence reconstruction agreement. For that, we used two measures: The Most Recent Common Ancestor (MRCA) (*12*), and The Correctness Of Ancestral Reconstruction (COAR) (*18*). The MRCA metric focuses on the most recent common ancestral, while COAR considers the entire evolutionary pathway. Both measures emphasize the importance of a correct ancestral reconstruction and do not penalize minor differences in the tree topologies whether the ancestral reconstruction is accurate.

The classical MRCA distance is calculated by iterating through all pairs of leaves. It compares the most recent ancestral sequence of each pair of leaves presenting into two trees simultaneously — for instance, the ground truth (*T*_1_) and the inferred tree (*T*_2_). Since internal nodes in the B cell lineage trees can also represent observed genotypes, we modified MRCA to consider all node pairs instead of leaves only. For a given pair of observed nodes *i* and *j*, where (*i j*,) ∈ *T*_1_ and (*i j*,) ∈ *T*_2_, the MRCA(*i j*,) is the normalized hamming distance between the most recent ancestral sequences in *T*_1_ and *T*_2_ trees, given by:

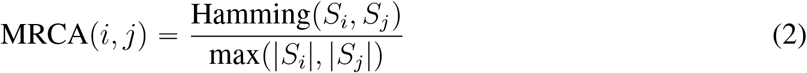

where *S*_*i*_ and *S*_*j*_ are the nucleotide sequences associated to nodes *i* and *j*, and Hamming(*S*_*i*_, *S*_*j*_,) computes the hamming distance between *S*_*i*_ and *S*_*j*_. Then, the MRCA metric is obtained by:

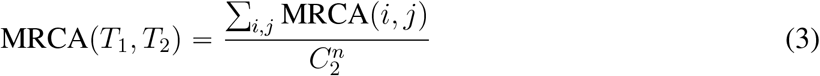

where *n* is the number of observable nodes in *T*_1_, or *T*_2_, and 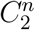 gives the total number of node pairs; see an example in Figure S3.

COAR (*12*) is another measure to evaluate the reconstruction of ancestral sequences. It compares evolutionary paths in the trees from the root (the naive sequence) to any leaf. To compute it, we consider each leaf *i* ∈ *T*_1_, find the path *p*_*i*_ from *i* until the root, compare it to all paths *p*_*j*_ ∈ *T*_2_ that contains *i*. To compare paths and obtain COAR(i), we used Needleman-Wunsch alignment algorithm with a score matrix based on negative hamming distance, and gap penalties. The COAR metric is the average of the COAR(i) of all leaves in *T*_1_:

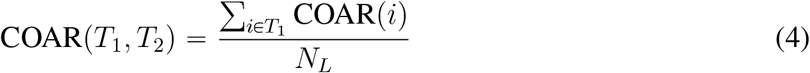

where *N*_*L*_ is the number of leaves in *T*_1_, and COAR(*i*) is computed by the Algorithm S1. See also, an example in Figure S3.

The numeric values of MRCA and COAR range in the interval [0, 1], where 0 represents a perfect ancestral sequence reconstruction, and 1 is the worst case. To see more details of these distance calculations, please refer to (*12*) and (*18*).

###### 3.2.1 Computational tools used for comparisons

There are a few B cell lineage reconstruction software; some are not available for download (*14*), others are specific for Windows operational system (*36*), and others are very timing consuming. Among the available methods, we selected two state-of-art tools to compare with ClonalTree: GCtree (*18*) and GLaMST (*21*). GCtree is an optimal solution that uses dnapars (*19*) to find a parsimony forest and then ranks parsimony trees according to genotype abundance information. On the other hand, GLaMST (*21*) is a time-efficient algorithm that uses MST but does not consider genotype quantities. The authors have shown a higher performance of these tools in the respective publications, GCTree outperformed IgPhyML (*12*), and GLaMST surpassed IgTree. Thus, we decided to keep only GCTree and GLaMST for evaluation and comparison.

## 4 Results and Discussion

We first validated our method with several artificial lineage trees that simulate affinity maturation. Then, we used two data sets containing experimental B cell lineages for biological validations.

### 4.0.1 Reconstructing B cell lineage trees from simulated data

To evaluate ClonalTree performance and compare it to GCtree and GLaMST, we generated simulated data sets using 92 different simulation settings, varying the root gene sequence and the relative probabilities of mutation, insertion, and deletion (see section 3.1 and Table 1). The artificial lineage trees served as ground truth that we would like to recover using B cell lineage tree algorithms. Thus, the performance measures how close reconstructed trees are to simulated ground truths. To quantify this resemblance, we used graph editing distances (GED) that measure the dissimilarity between two graphs/trees and two distances related to the correctness of common ancestral inferences. We computed two types of GED distances, based on the entire tree (GED tree-based) and separated paths (GED path-based), and used two previously defined metrics MRCA and COAR, to measure the correctness of ancestral reconstruction, see Section 3.2.

Figure 2 shows boxplots of GED distances for each compared method on the 92 simulated lineages trees. GCtree and ClonalTree had comparable performances, but ClonalTree outperformed GLaMST. Reconstructed B cell lineage trees of GCtree and ClonalTree displayed similar topologies, while trees produced by GLaMST are different. For GED tree-based distances (Figure 2-A), median values were 0, 2, and 12 and the highest distance 37, 38, and 120 for GCtree, ClonalTree and GLaMST, respectively. GLaMST presented the highest median value and the highest distance. ClonalTree produced 39 correct trees (GED tree-based distance equal to zero), while GLaMST produced only two trees.

**Figure 2:**
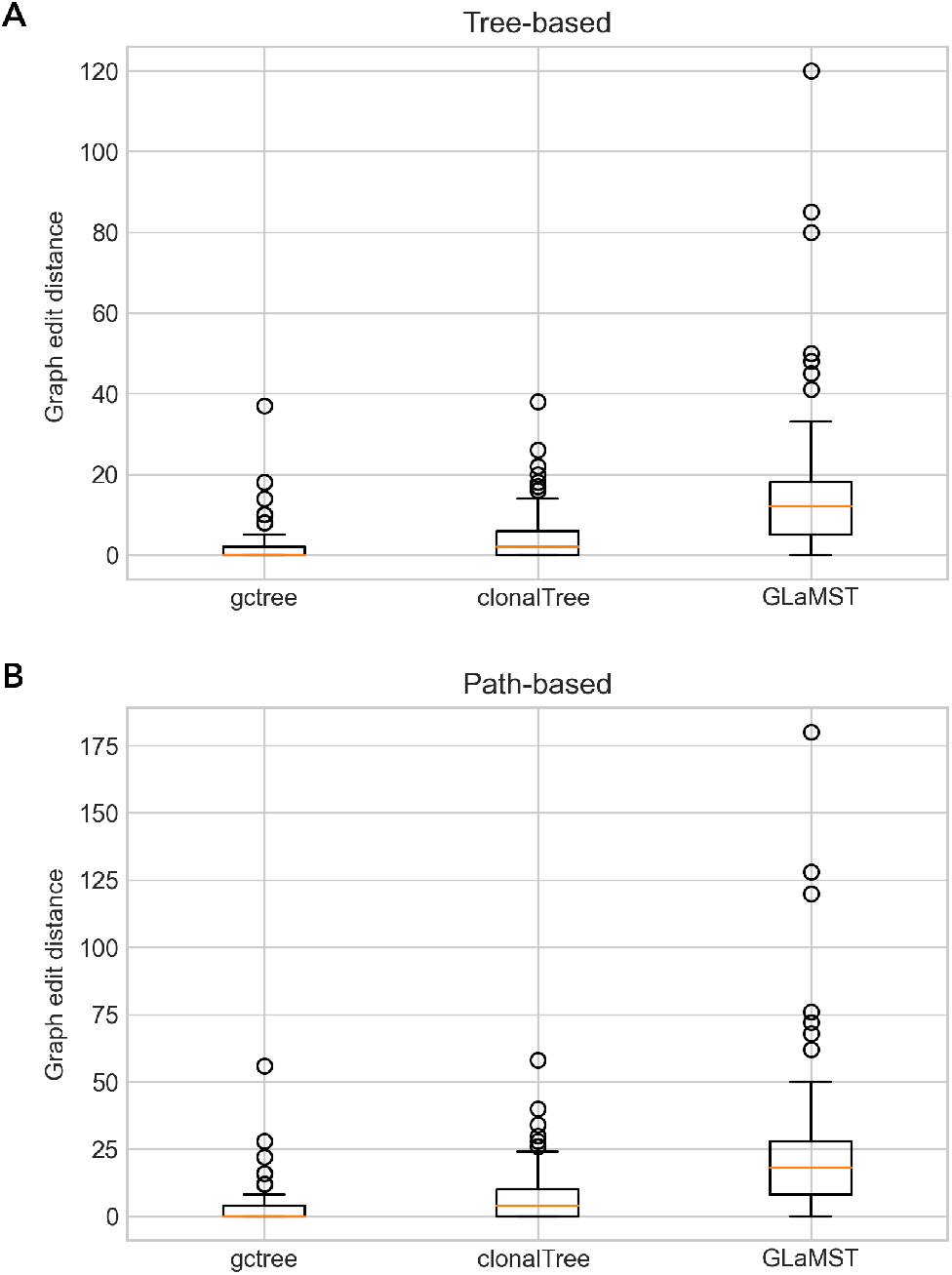
Performance comparison between GCtree, ClonalTree, and GLaMST using GED distances. Box-plots in (a) present GED tree-based distances, while (b) display GED path-based ones

GED path-based (Figure 2-B) compares each tree path, from leaves to the root, between inferred and ground truth trees; therefore, it is more sensitive to topology changes, resulting in higher distances than the GED tree-based ones. Figure 2-B confirms that GCtree and ClonalTree reconstructed B cell lineage trees with similar paths. Median values were 0, 4, and 18, with the highest distances 56, 58, and 180 for GCtree, ClonalTree, and GLaMST. As observed for GED tree-based, GLaMST presented the highest median value and distance. ClonalTree produced 39 correct paths (GED path-based distance equal to zero), while GLaMST produced only one path. In order to better evaluate the performance, we split the trees into three categories according to their number of sequences: small (between 30 and 50), medium (between 60 and 80), and large (having more than 90 sequences), see Referencesfig: percCat. We observed a slight difference between GCtree and ClonalTree, mainly in the small and large groups. On the other hand, we observed that GLaMST had difficulties in all groups; GED distances increased as the number of sequences grew.

The GED metric depends on the tree topologies, mainly GED-based-path that penalizes each path difference. Therefore, it is also essential to estimate the accuracy of ancestral reconstructions without penalizing minor differences in the tree topologies. For that, we used two metrics: the MRCA and the COAR, see Section 3.2. MRCA distance focuses on the most recent common ancestor and does not consider the entire evolutionary lineage. On the other hand, COAR measures the correctness of ancestral reconstruction from the root to any leaf. We first compared ClonalTree and GLaMST to ground truth trees and then with GCtree. The latter comparison is important since it gives us a basis for evaluating these methods on experimental data sets, where the true lineage evolution is unknown. Figure 3 shows MRCA distance distributions for ClonalTree and GLaMST when compared to ground truth trees (A) or GCtree (B). For both plots, we observed better performance for ClonalTree, which could reconstruct recent ancestral relationships more accurately. Figure 4 shows COAR distance distributions when comparing ClonalTree and GLaMST with ground truth trees (A) or with GCtree ones (B). Similarly, we observed that the trees produced by our approach are closer to both ground truth and GCtree trees. We noted that ClonalTree reconstructed trees that preserved the correctness of ancestral reconstruction, mainly when compared to ground truth trees.

**Figure 3:**
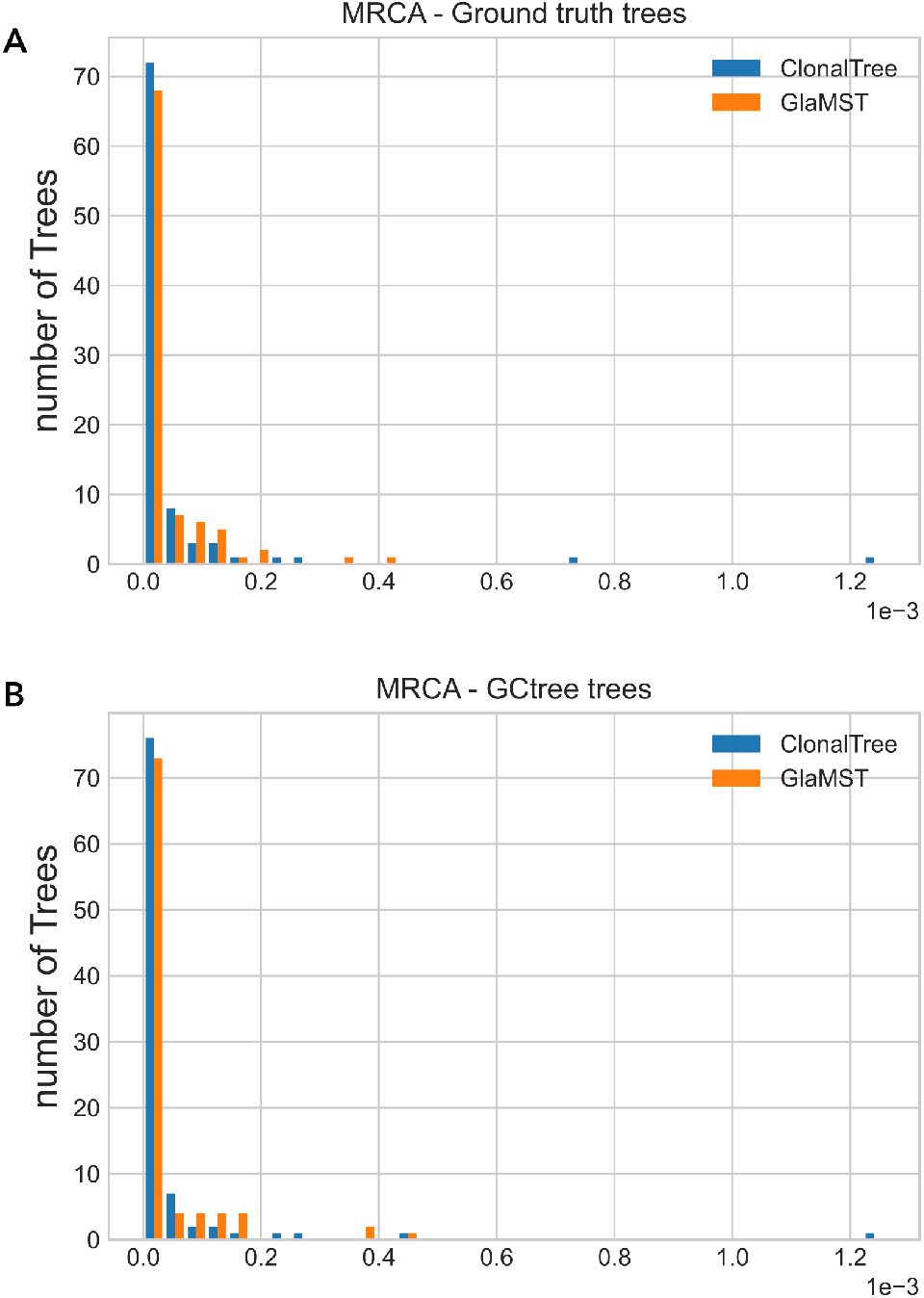
Most Recent Common Ancestor (MRCA) distance distributions. (a) compares Clonal-Tree, and GLaMST with ground truth trees, while (b) with trees generated by GCtree.

**Figure 4:**
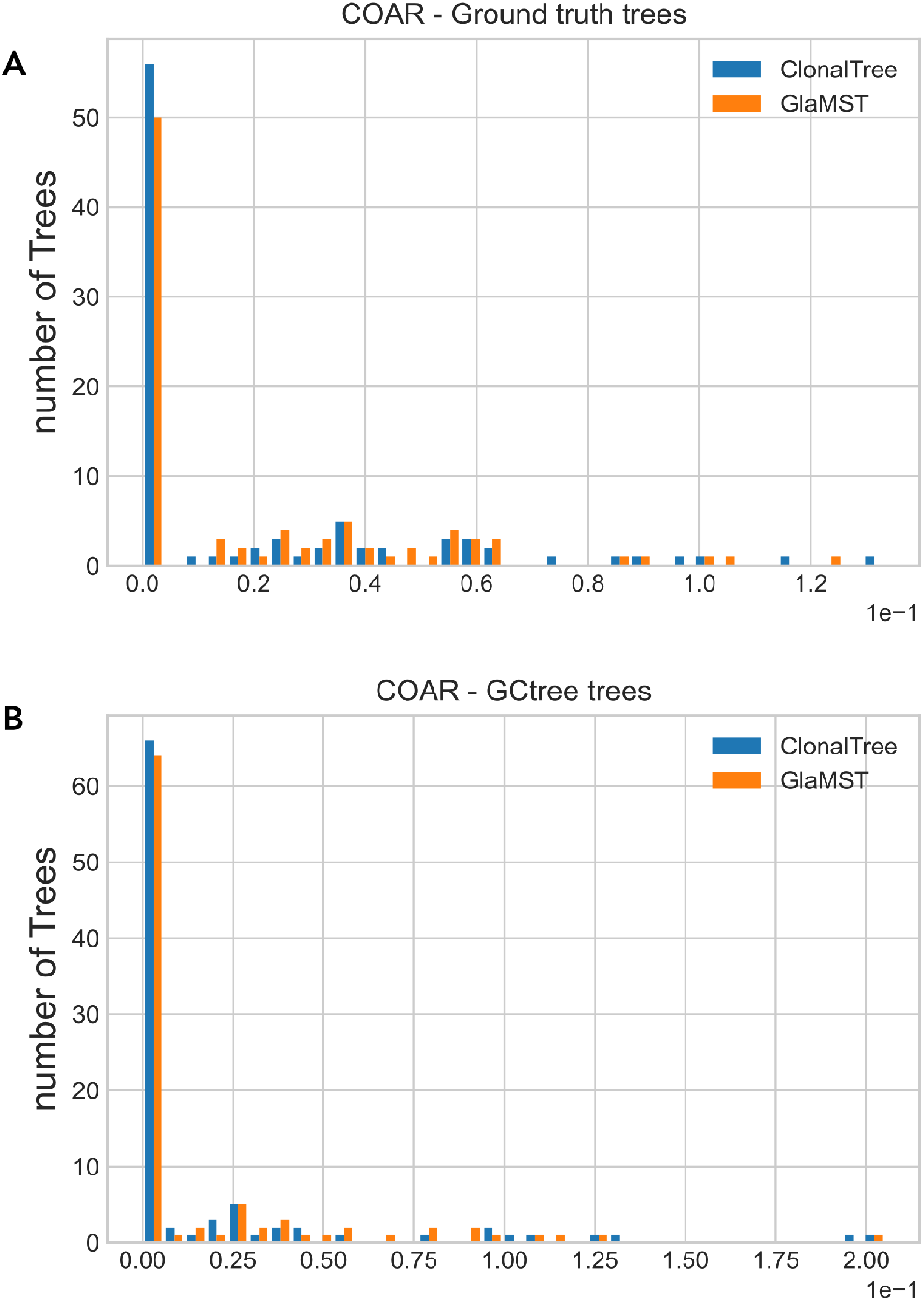
Correctness Of Ancestral Reconstruction (COAR) distance distributions. (a) compares ClonalTree, and GLaMST with ground truth trees, while (b) with trees generated by GCtree.

### 4.0.2 Biological validation using BCR sequencing data

We performed a biological validation on two experimental data sets: TAS-42 and CLL-20 (Section 3.1). Since ground truth trees are unavailable for these samples, we compared the inferred trees of ClonalTree and GLaMST with the trees inferred by GCtree. We considered trees generated with GCtree as references for the experimental validation since such a tool achieved the best performance on simulated data sets. The TAS-42 is a public data set generated by lineage tracing and single-cell germinal center BCR sequencing (*28*). TAS-42 contains 66 sequences, used as input for GCtree, GLaMST, and ClonalTree, see details in Section 3.1. The 2^*nd*^ and 3^*rd*^ columns of Table 2 show tree distances for ClonalTree and GLaMST when compared to GCtree, respectively. The GED tree-based distance was smaller for ClonalTree, producing a tree with ten differences from GCtree. GLaMST produced a more disparate tree than ClonalTree; Its GED tree-based distance was 47. GED path-based distances also showed that ClonalTree produced more similar evolutionary paths to GCtree than GLaMST. Only 22 evolutionary paths differ from GCtree against 230 for GLaMST. Most GlaMST paths contain various unobserved nodes, producing many mismatches compared to GCtree. We temporarily removed them to evaluate the tool’s performance. In the absence of unobserved nodes, ClonalTree presented just one mismatched path, while GLaMST produced seven. We also evaluated the agreement of ancestral sequence reconstructions; see Section 3.2. MRCA values were low for ClonalTree and GLaMST; both tools detected recent ancestral accurately for most node pairs. We did not observe a significant difference between MRCA values presented in Table 2. However, COAR distance values indicated that ClonalTree had better reconstructed entire evolutionary paths; we observed significantly lower COAR values for ClonalTree than GLaMST. Since ClonalTree achieved a lower GED path-based distance, it was expected to present lower COAR distances. COAR also considers sequence dissimilarities rather than only differences in the paths as GED path-based distance. Both measures confirmed that ClonalTree produces trees that are closer to GCtree ones.

**Table 2:** Performance evaluation of ClonalTree, and GLaMST on experimental BCR repertoire data sets. *GED path-based when removing unobserved nodes

| Metric | TAS-42 |  | CLL-20 |  |
| --- | --- | --- | --- | --- |
|  | ClonalTree | GLaMST | ClonalTree | GLaMST |
| GED tree-based | 10 | 47 | 3 | 36 |
| GED path-based | 22 | 230 | 18 | 1616 |
| GED path-based * | 1 | 7 | 2 | 13 |
| MRCA * $10^{-4}$ | 5.16 | 5.7 | 6.4 | 7.5 |
| COAR | 0.02 | 0.54 | 0.11 | 0.78 |

The CLL-20 data set was obtained from blood samples of a patient with CLL. We only analyzed the most abundant clone, which contains 3406 sequences and 20 different genotypes. For these sequences, IMGT/HighV-QUEST (*31*) inferred naive ancestor V(D)J genes using germline genetic references from IMGT (*37*) database, see Section 3.1. The 4^*th*^ and 5^*th*^ columns of Table 2 show tree distances for ClonalTree and GLaMST, respectively. ClonalTree GED tree-based distance was the smallest among experimental data sets, showing that ClonalTree generated a tree with a very similar topology to the one generated with GCtree. Likewise, GED path-based distance was smaller for ClonalTree. GLaMST presented an extremely high GED path-based distance; it added many unobserved nodes, causing several mismatches when comparing tree paths. We also evaluated tree path agreement without considering unobserved nodes. ClonalTree presented only two different paths against 13 of GLaMST. As observed in the data set TAS-42, we noted a slight difference between ClonalTree and GLaMST MRCA values, indicating that both tools reconstructed the most recent ancestral properly. However, a notable difference was again observed in COAR, being ClonalTree distance smaller than GLaMST one. This result is coherent with a better GED path-based distance since COAR accounts for the entire path from a leaf to the root.

The experimental results showed that the trees generated by ClonalTree were closer to GCtree ones; when compared to GLaMST, all distance metrics were smaller for ClonalTree, especially GED path-based and COAR distances. In summary, ClonalTree reconstructed more accurately the entire evolutionary lineage than GLaMST.

### 4.0.3 Time complexity and running time

The time complexity of a phylogenetic reconstruction algorithm is a function that represents the computing time required to analyze an instance of the problem. Typically, it depends on the number of treated sequences *n* and possibly other parameters. The computational cost of algorithms that exhaustively explore the tree generating space, producing eventually optimal solutions, increases considerably with input size. On the other hand, time-efficient approaches solve problems faster and more efficiently but can sacrifice optimality, accuracy, precision, or completeness. GCtree is exhaustive and finds the optimal solution, the most parsimonious tree. It uses dnapars with *O*(*n*^3^) complexity to produce a forest of equally parsimonious trees. Then GCtree ranks equally parsimonious trees through the abundance of genotypes. Although ranking parsimony trees requires just a polynomial increase in the dnapars runtime, finding the parsimony forest is computationally demanding and restricted to analyzing small-sized problems. MST-based algorithms are time-efficient; they are faster than GCtree because they construct a single tree instead of a forest. Their time complexity is determined by the MST algorithm’s complexity and the pairwise distance calculation. GLaMST and ClonalTree use Prim’s algorithm to grow the tree from the root node and interactively add edges and nodes (possibly unobserved). The time complexity of Prim’s algorithm is *O*(|*E*| + log |*V* |), where |*V* | = *n* and *E* is the number of edges or connections among nodes. However, GLaMST computes pairwise edit distances using a dynamic programming algorithm with time complexity around *O*(*n*^2^). Contrarily to GCtree and GLaMST, ClonalTree uses hamming distance since sequences are previously aligned with Clustal Omega (*38*). Both alternatives decrease the ClonalTree time complexity substantially. Hamming distance is computed in *O*(*n*), while multiple sequence alignment in *O*(*n* * (log(*n*))^2^). In order to measure the running time of different tools, we used the data set provided by GLaMST, containing 684 observed sequences. The three algorithms were compared on a desktop workstation with i7 processor at 3.40GHz and 32G memory. GCtree took 4.5 days, GLaMST spent 55 minutes, while ClonalTree took less than 15 minutes to reconstruct a lineage tree for this data set. Note that we used a pre-compiled version of GLaMST that is slower than the source code. GLaMST was coded in Matlab, a proprietary programming language, and only an executable for Windows is available for users that do not possess a Matlab license. Since ClonalTree is coded in python and uses freely available libraries and software, it is ready for the scientific community.

## 5 Conclusion

We addressed the computational problem of reconstructing lineage trees from high-throughput BCR sequencing data. It is challenging since this data often contains a subset of BCR IGH sequences, partially representing the dynamic process of BCR affinity maturation. Therefore, efficient algorithms must reconstruct the entire evolutionary history from partially observed sequences. We proposed ClonalTree, a fast method that combines minimum spanning trees with genotype abundance information to reconstruct accurate evolutionary trees. ClonalTree outperformed GLaMST, a method based only on the minimum spanning tree approach, on simulated and experimental data. ClonalTree has systematically produced more maximum parsimonious trees than GLaMST.

Compared to GCtree, an exhaustive sequence-based phylogenetic method, ClonalTree presented a comparable performance, mainly on experimental data, where we observed few differences in the produced trees. However, ClonalTree is computationally more efficient, achieving equivalent results with a lower runtime. ClonalTree was hundreds to thousands of times faster than GCtree, allowing the analysis of large data sets within minutes or seconds and with minor loss of accuracy. ClonalTree’s high and fast performance allows the users to consider all the available genotypes when reconstructing lineage trees. It can help researchers understand B cell receptor affinity maturation, mainly when data from a dense quantitative sampling of diversifying loci are available. Integrating ClonalTree into existing BCR sequencing analysis frameworks could speed up lineage tree reconstructions without compromising the quality of evolutionary trees.

## Acknowledgements

Authors are grateful to Thibaud Verny for his insightful comments and fruitful discussion. We also acknowledge the anonymous reviewers that contributed to improve the manuscript and the method.

## Supporting information

**Figure S1** Editing the reconstructed B cell lineage tree by adding unobserved internal nodes

**Figure S2** Editing the reconstructed BCR lineage tree to reduce the depth of the tree while keeping its overall

**Figure S3** An example of MRCA calculation.

**Figure S4** An example of COAR calculation for a given leaf.

**Figure S5** Performance comparison among GCtree, ClonalTree, and GLaMST using GED distances on three

**Algorithm S1** COAR algorithm.

