## Supplementary Figures for "Reconstructing the evolutionary history of a B cell lineage with minimum spanning tree and genotype abundances"

September 30, 2022

### Supplementary Figures

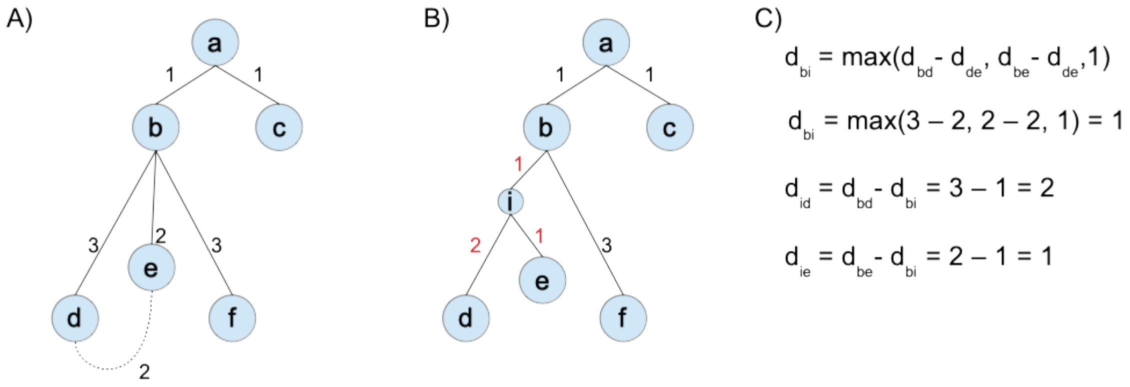


Figure 1: **Editing the reconstructed B cell lineage tree by adding unobserved internal nodes**. Unobserved internal nodes might represent unobserved sequences that were not sampled or disappeared during the affinity maturation When the distance between two sister nodes is smaller or equal to the distance to their parent (dotted line in A), we add an unobserved internal node as the common ancestor of the two sister nodes (node i in B). C shows how to update edge weights (in red).

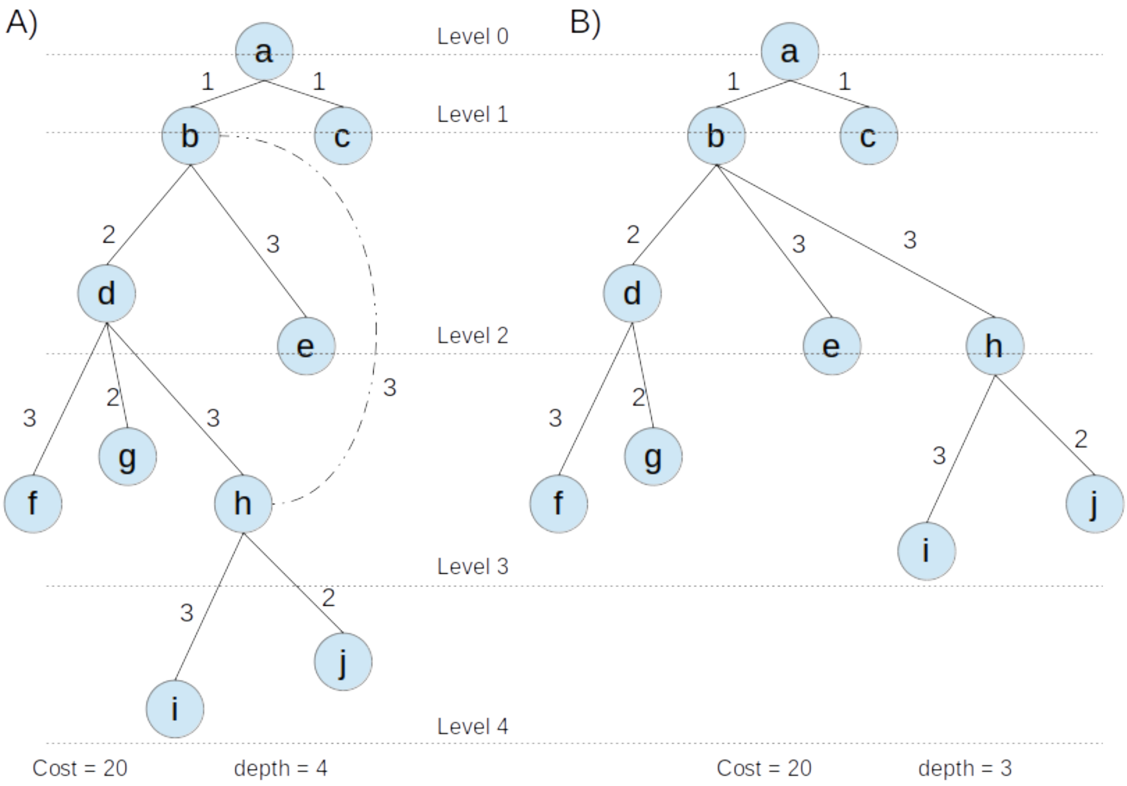


Figure 2: **Editing the reconstructed B cell lineage tree to reduce the depth of the tree while keeping its overall cost.** In this example, the total cost of the tree, or the sum of edge weights, remained the same while the depth of it has been reduced. Edge weights represent the Hamming distance between sequences.

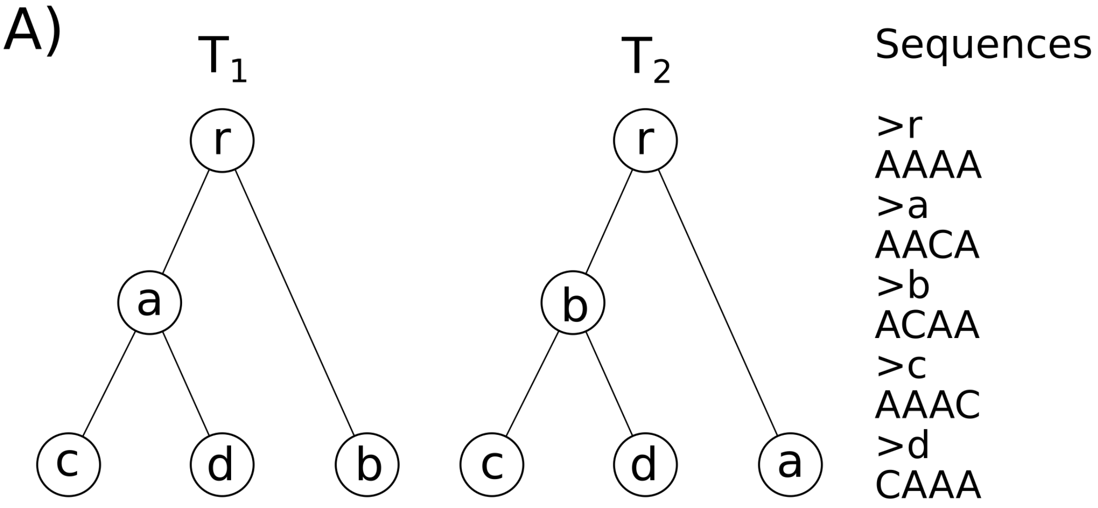


B)

| Node pair(i,j) | Ancestral in $T_1$ | Ancestral in $T_2$ | MRCA(i,j) |
|---|---|---|---|
| c, d | a | b | 2/4 |
| c, a | a | r | 1/4 |
| c, r | r | r | 0 |
| c, b | r | b | 1/4 |
| d, a | a | r | 1/4 |
| d, b | r | b | 1/4 |
| d, r | r | r | 0 |
| a, r | r | r | 0 |
| a, b | r | r | 0 |
| b, r | r | r | 0 |
| | | $\sum MRCA(i,j)$ | 6/4 |

C) $$MRCA(T_1, T_2) = \frac{\sum MRCA(i,j)}{C_2^5} = \frac{6/4}{10} = 0.15$$

Figure 3: **An example of MRCA calculation.** A) Let's $T_1$ and $T_2$ be two comparable trees, and Sequences a fasta file containing nucleotide sequences associated to each node in both trees. B) For each pair of nodes $(i,j) \in T_1$ and $(i,j) \in T_2$, we find its most recent ancestral in both tree, and computed MRCA(i,j) as the normalized hamming distance bteween ancestral sequences. C) MRCA($T_1, T_2$) is the average of MRCA(i,j) of all pair of nodes.

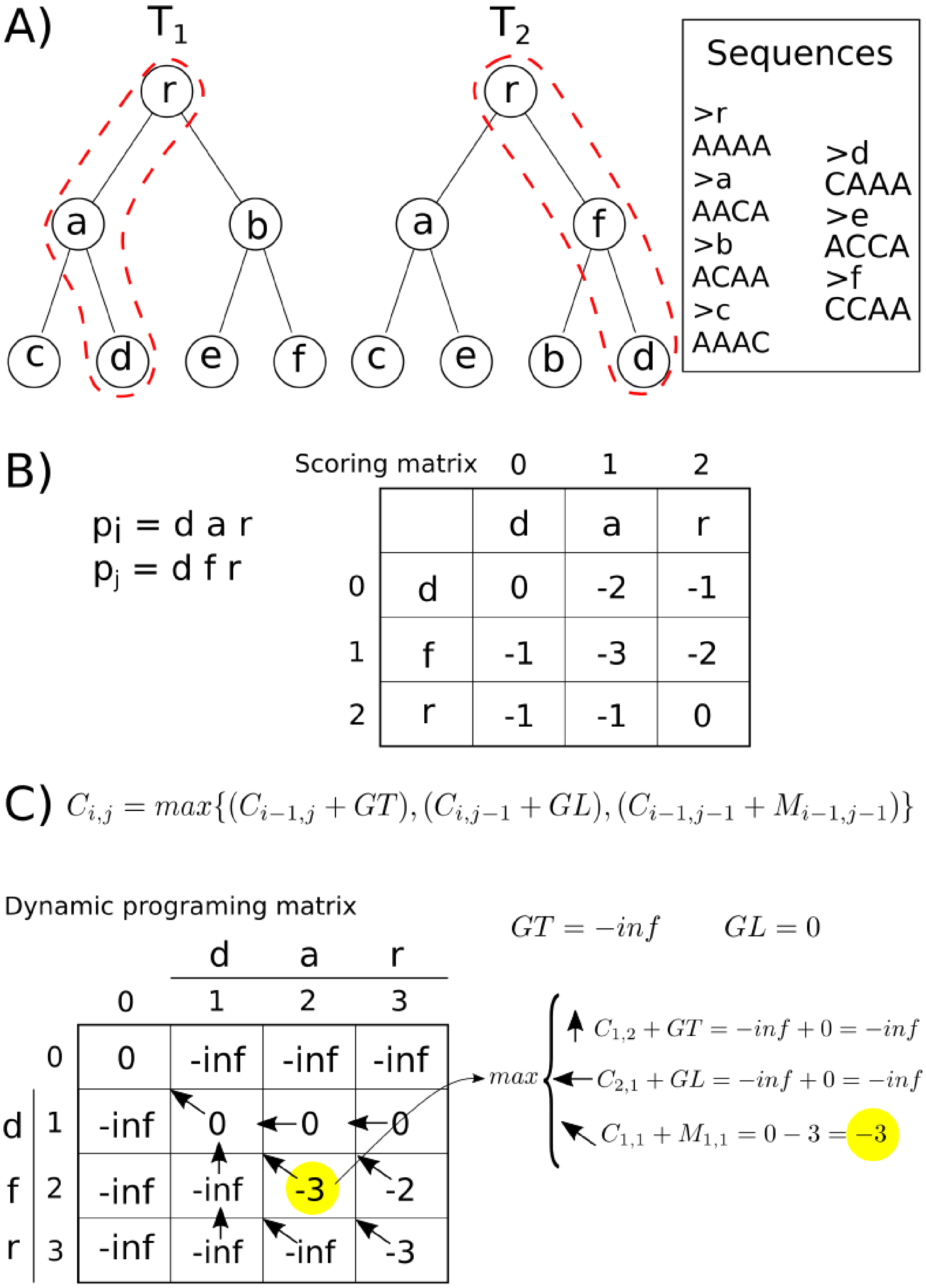


Figure 4: **An example of COAR calculation for a given leaf (see the algorithm below)** A) Let's $T_1$ and $T_2$ be two comparable trees, and Sequences a fasta file containing nucleotide sequences associated to each node in both trees. Let's dotted paths $p_i$ and $p_j$ represent comparable paths in $T_1$, and $T_2$, respectively. B) Paths to be compared and the scoring matrix containing negative hamming distances for each pair of sequence $S_i \in p_i$ and $S_j \in p_j$. C) We align paths with Needleman-Wunsch alignment algorithm, by using the scoring matrix computed in B, and special gap penalties: -inf for gap top (GT), and 0 for gap left (GL). Each element of the dynamic programming matrix is computed by the formula $C_{i,j}$, see an example for cell $C_{2,2}$. The COAR for the leaf $i$ is $1 = \frac{minScore_i}{min(M)} = \frac{C_{3,3}}{-3} = \frac{-3}{-3}$.

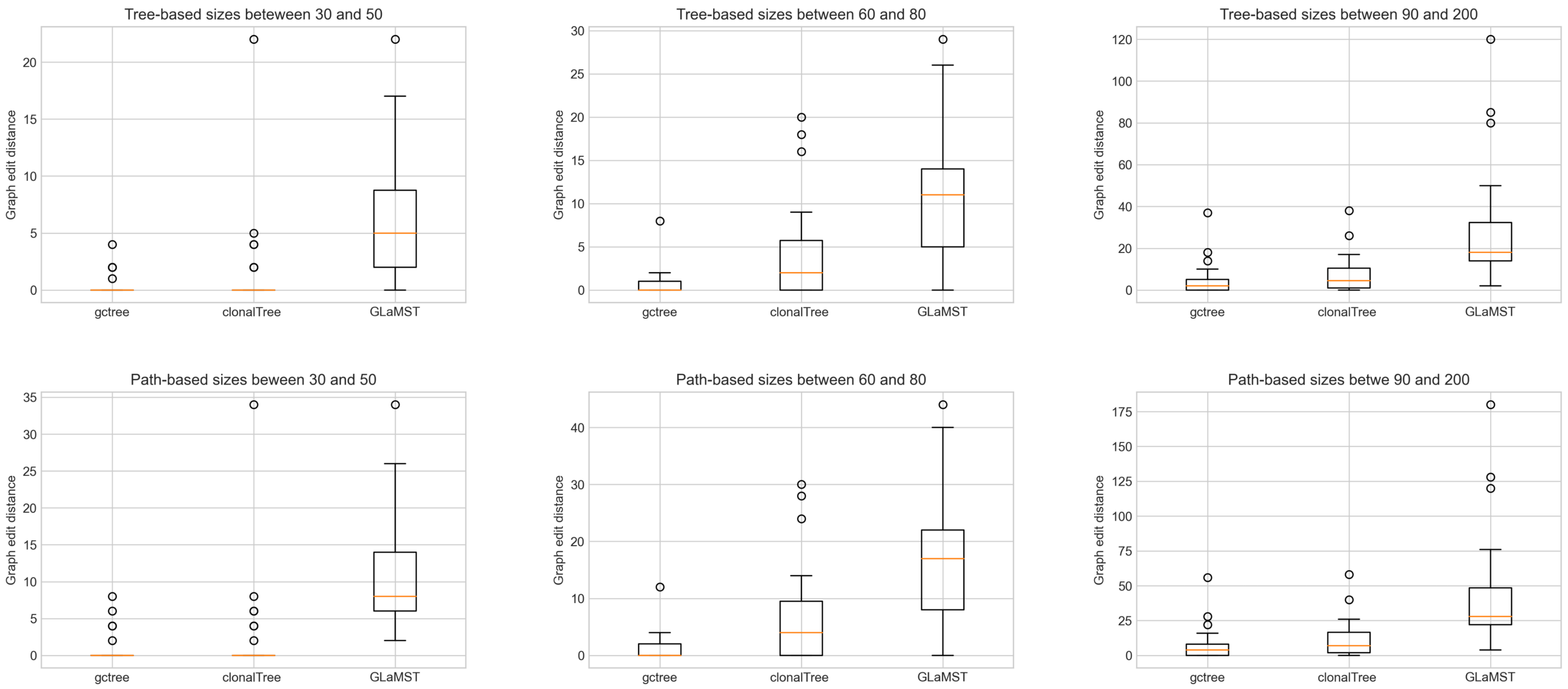


Figure 5: **Performance comparison among GCtree, ClonalTree, and GLaMST using GED distances on three categories of trees.** The categories are based on the tree sizes.

**Algorithm 1:** COAR algorithm

**Require:** $T_1, T_2, \text{sequences}, \text{GapPenalties}$
  $COAR \leftarrow 0$
  $N_L \leftarrow 0$ {Number of Leaves}
  **for all** leaf $i \in T_1$ **do**
    $N_L \leftarrow N_L + 1$
    $p_i \leftarrow path(i, T_1)$
    $P \leftarrow paths(i, T_2)$ {take all paths in $T_2$ containing i}
    $minScore_i \leftarrow inf$
    {Compute a scoring matrix for sequences associated to nodes in $p_i \bigcup P$ based on negative hamming distances}
    $M \leftarrow scoreMatrix(P, p_i, sequences)$
    **for all** $p_j \in P$ **do**
      $scoreAln_i \leftarrow NWS(p_i, p_j, M, GapPenalties)$
      **if** $scoreAln_i > minScore_i$ **then**
        $minScore_i \leftarrow scoreAln_i$
      **end if**
    **end for**
    $COAR_i \leftarrow \frac{minScore_i}{min(M)}$
    $COAR \leftarrow COAR + COAR_i$
  **end for**
  **return** $\frac{COAR}{N_L}$

COAR algorithm requires two comparable trees $T_1$ and $T_2$, a set of nucleotide sequences, and Gap penalties. For each leaf $i \in T_1$, we find its path until the root, named $p_i$. We also compute $P$, a list of paths in $T_2$, containing $i$. Note that if $i$ is a leaf in $T_2$, then $|P| = 1$, otherwise $|P| > 1$. We then compute a scoring matrix $M$ for all nodes in $p_i \bigcup P$, each element of $M$ contains the negative hamming distance between nodes' representative sequences. For instance, let a and b be nodes, and $S_a$='ACCA' and $S_b$='CCCC' the nucleotide sequences associated with those nodes. Then $M_{a,b}$ = -2. Next, for each $p_j \in P$, we align it with $p_i$ to obtain the scoreAln$_i$. For that, we use the Needleman-Wunsch (NWS) algorithm, requiring the scoring matrix, previously computed, and Gap Penalties. In order to avoid gaps in the longest path, we use an NWS version that puts the longest sequence in the columns of the dynamic programming matrix and uses different gap penalties: 0 for the gap left (GL) and -inf for the gap top (GT). In case of $|P| > 1$, we keep the minimum alignment score for $i$. Finally, we compute COAR$_i$ as the ratio between minScore$_i$ and the minimum value in M. We iterate until we compute COAR$_i$ for all $i \in T_1$, and we return the average: the overall COAR divided by the number of leaves in $T_1$. We ilustrate a iteration of COAR algorithm above.
